# Arginine impacts aggregation, biofilm formation, and antibiotic susceptibility in *Enterococcus faecalis*

**DOI:** 10.1101/2024.05.30.596650

**Authors:** Alex Snell, Dawn A. Manias, Reham R. Elbehery, Gary M. Dunny, Julia L. E. Willett

## Abstract

*Enterococcus faecalis* is a commensal bacterium in the gastrointestinal tract (GIT) of humans and other organisms. *E. faecalis* also causes infections in root canals, wounds, the urinary tract, and on heart valves. *E. faecalis* metabolizes arginine through the arginine deiminase (ADI) pathway, which converts arginine to ornithine and releases ATP, ammonia, and CO_2_. *E. faecalis* arginine metabolism also affects virulence of other pathogens during co-culture. *E. faecalis* may encounter elevated levels of arginine in the GIT or the oral cavity, where arginine is used as a dental therapeutic. Little is known about how *E. faecalis* responds to growth in arginine in the absence of other bacteria. To address this, we used RNAseq and additional assays to measure growth, gene expression, and biofilm formation in *E. faecalis* OG1RF grown in arginine. We demonstrate that arginine decreases *E. faecalis* biofilm production and causes widespread differential expression of genes related to metabolism, quorum sensing, and polysaccharide synthesis. Growth in arginine also increases aggregation of *E. faecalis* and promotes decreased susceptibility to the antibiotics ampicillin and ceftriaxone. This work provides a platform for understanding of how the presence of arginine in biological niches affects *E. faecalis* physiology and virulence of surrounding microbes.

## Introduction

*Enterococcus faecalis* is found as a low-abundance commensal bacterium in the gastrointestinal (GI) tract of humans, animals, and insects (1, 2). *E. faecalis* can be highly resistant to antibiotics and other antimicrobial treatments, and it forms robust biofilms on surfaces (3, 4). Outside of the GI tract, *E. faecalis* is a prevalent cause of opportunistic infections in wounds, heart valves, implanted devices, the urinary tract, and root canals (4–6). In these environments, *E. faecalis* interacts with other microbes and promotes virulence of pathogens including *Clostridioides difficile*, uropathogenic and enterohemorrhagic *Escherichia coli*, *Pseudomonas aeruginosa*, *Proteus mirabilis*, and *Staphylococcus aureus* (7–13). A key nutrient mediating many of these interactions is arginine. *E. faecalis*, like many bacteria, can metabolize arginine as an energy source through the arginine deiminase (ADI) pathway, producing ATP, L- ornithine, CO_2_, and NH_3_ as byproducts (14, 15). Export of L-ornithine by *E. faecalis* promotes siderophore synthesis by *E. coli*, and breakdown of arginine by *E. faecalis* promotes virulence of *C. difficile* (10, 13). Disrupting the ADI pathway can also affect *E. faecalis* biofilm formation *in vitro* (16, 17).

The ADI pathway is relevant to health and disease in the oral cavity (18). Production of NH_3_ by the oral microbiome and the resulting increase in pH can protect against acidification caused by bacteria such as *Streptococcus mutans*, as low pH can lead to the development of caries (cavities) (19). L-arginine reduces biofilm formation independent of growth inhibition in a multi-species biofilm with oral bacteria and led to growth suppression of caries-causing *S. mutans* in biofilms with *Streptococcus gordonii* and *Actinomyces naeslundii* (20, 21).

Application of exogenous arginine results in destabilization and disruption of pre-formed biofilms grown from *S. gordonii* monocultures or a mixed oral bacterial community (20, 22). In patients with caries, the use of toothpaste supplemented with arginine led to an increase in ADI pathway activity and an increase in plaque species that are associated with health instead of disease (23). The ADI pathway also affects biofilms and antibiotic tolerance of *Streptococcus pyogenes* (24), and exogenous arginine decreases biofilm formation in *Staphylococcus aureus* (25). Despite the importance of arginine catabolism and ADI pathway byproducts on biofilm formation and polymicrobial interactions involving numerous Gram-positive bacteria, little is known about the effect of arginine on gene expression, biofilm formation, and antimicrobial resistance in *E faecalis*.

Our goal was to determine how arginine affects *E. faecalis* independent of polymicrobial interactions. Here, we used *E. faecalis* OG1RF to measure changes in gene expression and biofilm formation after growth in arginine relative to growth in control medium without arginine. We found that similar to other bacteria, growth in arginine decreased biofilm formation by *E. faecalis*. However, pre-formed *E. faecalis* biofilms were not susceptible to destabilization by high concentrations of arginine as has been reported for oral streptococci. We also found widespread changes in gene expression in arginine-grown cells relative to control cells, including changes in expression of metabolic pathways and virulence factors. Growth in arginine promoted aggregation, increased cell envelope permeability, and increased resistance to select cell wall-targeting antibiotics, including ampicillin. Together, this demonstrates that the presence of arginine induces global changes in the growth and physiology of *E. faecalis*, which will serve as the basis for future studies on how these changes affect monomicrobial and polymicrobial infections involving *E. faecalis*.

## Results

### Arginine promotes aggregation and decreases biofilm formation of *E. faecalis*

In *E. faecalis* and related bacteria, arginine is metabolized through the arginine deiminase (ADI) pathway, which generates ATP and leads to a concomitant increase in optical density relative to un-supplemented cultures (14). To confirm this in our system, we used the well-studied strain OG1RF, which has been used in other studies demonstrating that arginine metabolism and ornithine production enhance virulence and polymicrobial infections (10, 13, 26). We grew OG1RF in 96-well plates using complex basal medium (CBM) supplemented with arginine (25 or 50 mM) and measured A_600_ over time. CBM was chosen as it was originally used to study arginine as an energy source in *E. faecalis* (14). Untreated CBM or CBM supplemented with glucose (50 mM) or glycine (25 or 50 mM) were used as controls. These concentrations were chosen as they have been previously used to study the ADI pathway in *E. faecalis* and biofilm formation and dispersal in oral streptococci (14, 15, 27). For cultures grown with arginine, the A_600_ increased relative to untreated CBM, but the final A_600_ was not as high as glucose-grown cultures (**Figure 1A**). No increase was observed with glycine. We next grew 3 mL cultures in glass tubes for 24 hr and quantified CFU/mL. As expected, the CFU/mL of cultures grown with glucose was ∼0.5 log higher than untreated CBM cultures, but only a small increase (∼0.1 log) was observed with arginine supplementation (**Figure 1B**). Similarly, the A_600_ of glucose-grown cultures in test tubes was higher than untreated CBM cultures, but arginine did not lead to a consistent increase in the A_600_ in these cultures like in 96-well plates (**Figure 1C**).

**Figure 1.**
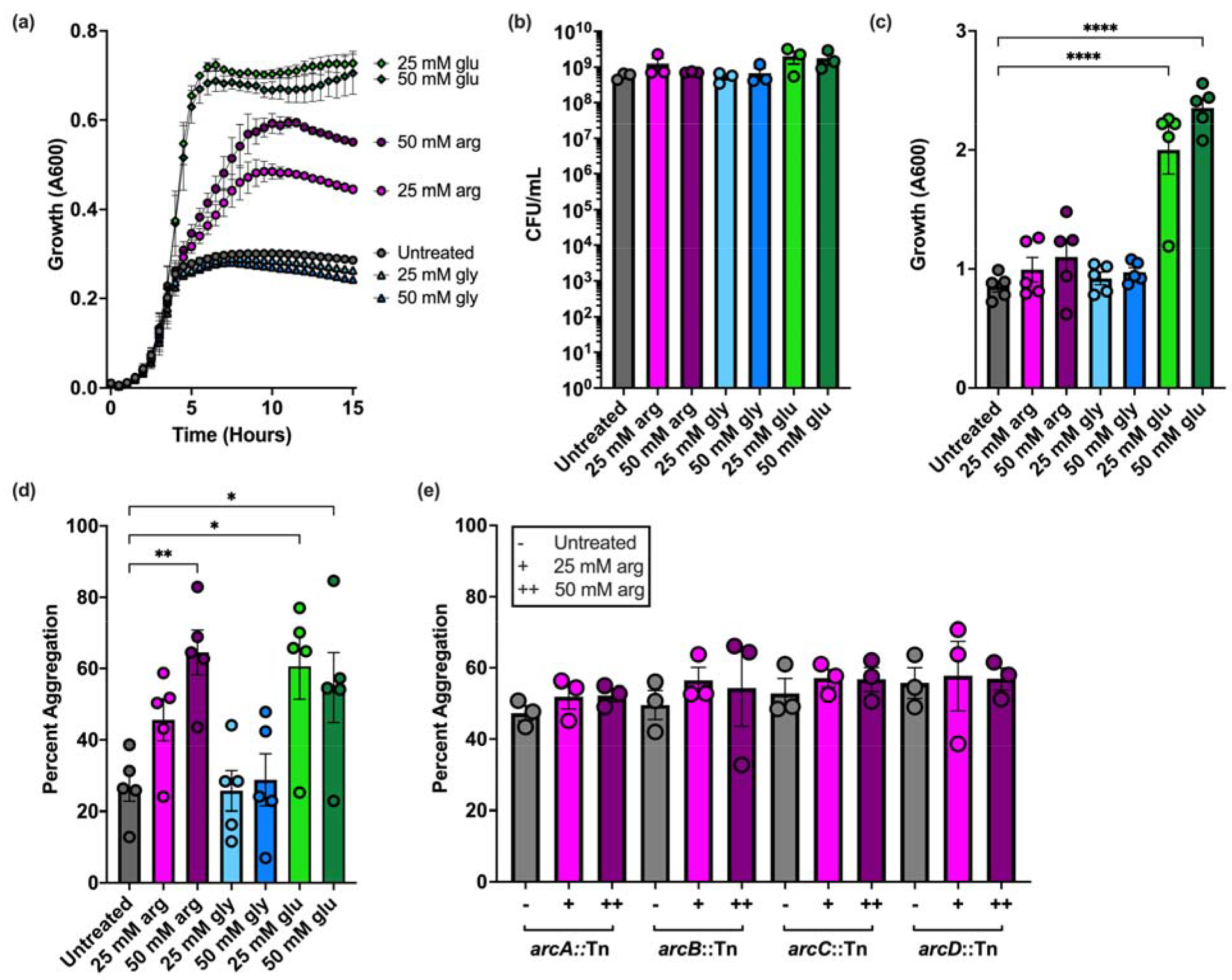
Growth and aggregation of *E. faecalis* OG1RF grown in arginine. (a) Microtiter plate growth curve of OG1RF grown in CBM supplemented with the indicated concentrations of arginine, glycine, or glucose. **(b)** CFU/mL of OG1RF grown in glass test tubes in CBM supplemented with arginine, glycine, or glucose for 24 hr. **(c)** Absorbance at 600 nm of OG1RF grown in glass test tubes in CBM supplemented with arginine, glycine, or glucose for 24 hr. **(d)** Aggregation assay of OG1RF grown with arginine, glycine, and glucose for 24 hr. **(e)** Aggregation of Tn mutants with insertions in genes encoding the ADI pathway and an arginine/ornithine antiporter (*arcABCD*) after growth in untreated CBM or arginine for 24 hr. In panel **(a)**, data points represent the average of three independent biological replicates. For panels **(b)**-**(e)**, each data point represents one independent biological replicate with 2 or 3 technical replicates. For all panels, error bars represent standard error of the mean. Statistical significance in panels **(c)** and **(d)** was determined by one-way ANOVA with Tukey’s test for multiple comparisons. Comparisons that were not statistically significant are not shown. *p<0.05, **p<0.01, ***p<0.001, ****p<0.0001.

During these experiments, we observed aggregation of cells at the bottom of culture tubes and quantified the percent of aggregated cells in each condition. In 50 mM arginine, the percent of aggregated cells rose to ∼60% on average relative to ∼25% in untreated CBM (**Figure 1D**).

Aggregation also increased in glucose-grown cells to a similar level as arginine-treated cultures, but no increase was observed with glycine. Arginine-dependent aggregation was not observed in Tn mutants with insertions in the ADI pathway genes *arcABC* or *arcD*, which encodes an arginine/ornithine antiporter (**Figure 1E**). Together, these results suggest that catabolism of arginine by *E. faecalis* leads to an increase in aggregation.

Next, we asked how arginine affected *E. faecalis* biofilm formation, given that arginine reduces biofilm formation in other Gram-positive bacteria (24, 25, 27). We grew OG1RF biofilms for 6 or 24 hr in 96-well plates using the media described above and quantified safranin-stained biofilm material (A_450_) relative to A_600_. In early biofilms (6 hr), growth in 25 and 50 mM arginine reduced OG1RF biofilm by ∼50% relative to untreated CBM, but glycine did not affect the level of biofilm produced (**Figure 2A**). At 24 hr, biofilm formation in arginine-supplemented cultures was reduced 60-75% relative to untreated biofilms and those grown with glycine (**Figure 2B**). We next asked whether exogenous arginine added after biofilm formation could disrupt pre-formed biofilms. Biofilms were grown for 6 or 24 hr, after which arginine, glycine, or a buffer control was added for 20 min. No dispersal or decrease in biofilm level relative to control biofilms was observed. Surprisingly, post-growth treatment of early biofilms with glycine resulted in an increase in biofilm index values relative to control-treated biofilms.

**Figure 2.**
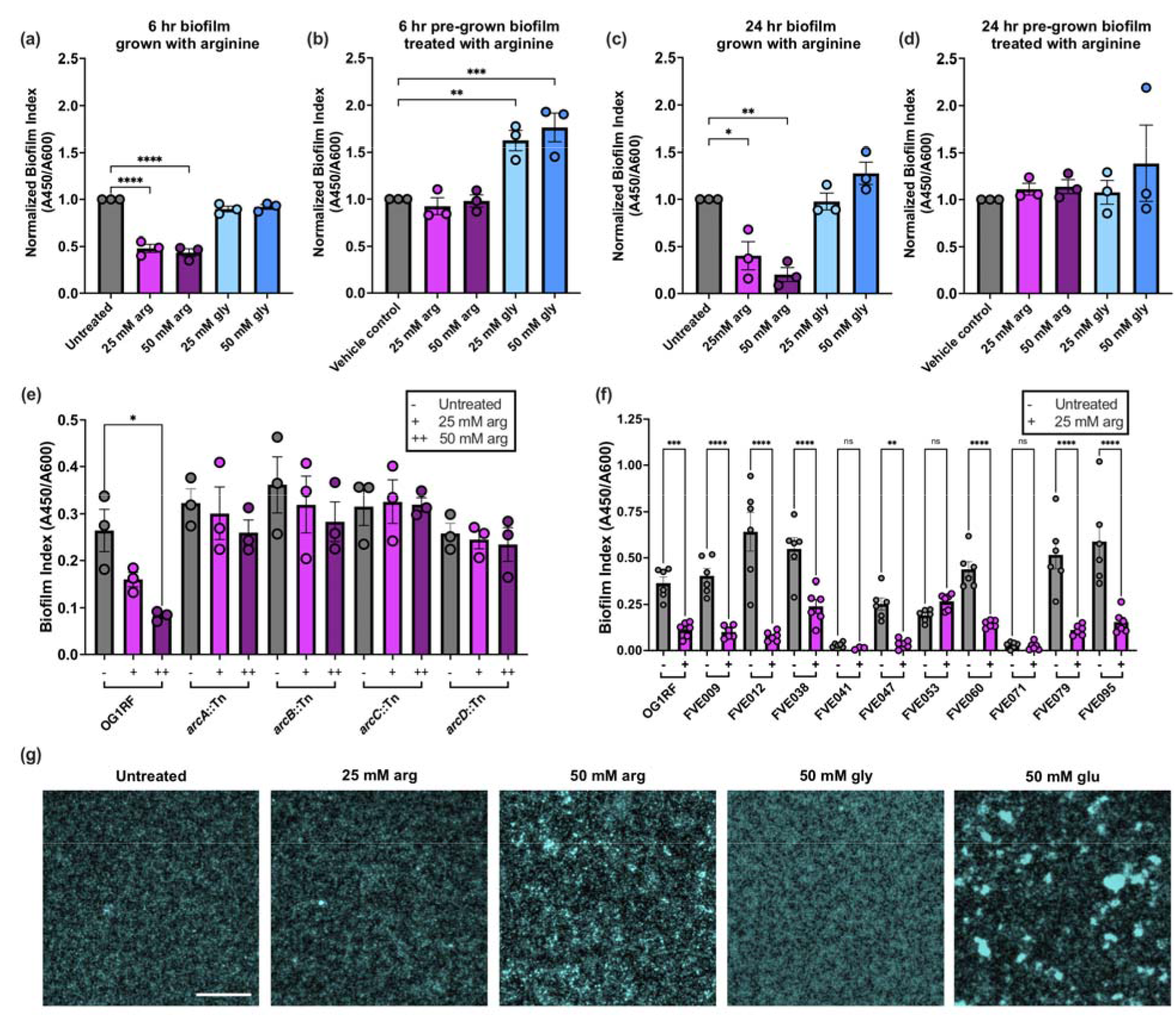
Arginine affects biofilm formation and structure in *E. faecalis*. Biofilm formation was measured in microtiter plates with arginine added **(a)** prior to growth for 6 hr, **(b)** after 6 hr growth, **(c)** prior to growth for 24 hr, and **(d)** after 24 hr growth. For pre-formed biofilms, arginine was added for 20 min and removed by washing. Biofilm index values were normalized to those measured in un- supplemented CBM or with a vehicle control. **(e)** 24 hr microtiter plate biofilm formation of OG1RF and Tn mutants with insertions in ADI pathway genes *arcABCD*. **(f)** 24 hr microtiter plate biofilm formation of *E. faecalis* gastrointestinal tract isolates from healthy human volunteers. **(g)** Fluorescence microscopy visualization of *E. faecalis* OG1RF biofilms grown for 24 hr in the indicated conditions. Cells were fixed in formalin and stained with Hoechst 33342. The scale bar represents 50 μm. Images shown are representative of three independent biological replicates. For panels **(a)-(f)**, each data point represents one independent biological replicate with three technical replicates. Error bars represent standard error of the mean. Statistical significance was determined by one-way ANOVA with Tukey’s test for multiple comparisons. Comparisons that were not statistically significant are not shown. *p<0.05, **p<0.01, ***p<0.001, ****p<0.0001

We speculate that this may be due to glycine binding to or interacting with a component of the early biofilm matrix, since this increase was not evident at 24 hr. We next measured biofilm formation and destabilization in higher concentrations of arginine (100, 250, and 500 mM) that have been used therapeutically and for other studies on oral biofilms (18, 20, 21, 28, 29). We observed a 70-95% reduction in biofilm formation when arginine was added during growth, but surprisingly, addition of 500 mM exogenous arginine did not significantly destabilize pre-formed biofilms (**Supplementary** Figure 1), even though this concentration was shown to disrupt oral biofilms *in vitro* (20, 21, 27).

To test whether this disruption in biofilm formation was dependent on arginine metabolism through the ADI pathway, we cultured mutants with Tn insertions in *arcABCD* and measured biofilm formation with and without arginine. An arginine-dependent decrease in biofilm formation was not observed in the *arcABCD* Tn mutants (**Figure 2E**). Interestingly, we previously observed a reduction in biofilm formation relative to parental OG1RF with these mutants in tryptic soy broth without added glucose (17), suggesting that other components of growth medium may influence biofilm formation via the ADI pathway. To ensure that this response was not specific to OG1RF, we measured microtiter plate biofilm formation after 24 hr growth in the presence and absence of arginine for 10 *E. faecalis* isolates obtained from fecal samples from healthy human volunteers (30). Arginine-dependent decreases in biofilm formation were observed for 7 of the 10 strains (**Figure 2F**). Two of the strains that did not have reduced biofilm formation in the presence of arginine formed very poor biofilms in control conditions. Therefore, we conclude that arginine disruption of *E. faecalis* biofilm formation is not a strain-specific effect.

Finally, we asked how biofilm morphology was impacted by growth in arginine and used fluorescence microscopy to evaluate the appearance of biofilms grown for 24 hr. OG1RF grown in un-supplemented CBM formed monolayer biofilms (**Figure 2G**), similar in appearance to those observed with OG1RF (31–33). Biofilms grown with 25 mM arginine resembled those grown in CBM, but growth in 50 mM arginine led to the formation of biofilms with aggregates of cells. Growth in glycine resulted in monolayer biofilms similar in appearance to those grown in CBM, whereas growth in glucose resulted in biofilms with large multi-cellular aggregates, as has been previously observed in glucose-rich media (32). This data demonstrates that growth in arginine alters *E. faecalis* biofilm formation via the ADI pathway.

### Transcriptional response of *E. faecalis* to arginine

Because we observed significant changes in aggregation and biofilm formation after growth in arginine, we sought to determine how arginine affects global gene expression in *E. faecalis*. We grew planktonic and biofilm cultures of OG1RF in CBM in the presence and absence of 25 mM arginine for 6 hr and used RNAseq to compare differentially expressed transcripts (**Supplementary Table 1**, **Figure 3**). Relatively few loci (genes or intergenic regions) were differentially expressed in planktonic compared to biofilm culture relative to the number of genes differentially expressed in the presence of arginine. 34 loci were downregulated in biofilms relative to planktonic culture in the absence of arginine, and 8 were upregulated. 35 loci were downregulated in biofilms relative to planktonic culture in arginine, and 8 were upregulated. In planktonic culture, 1037 loci were downregulated and 909 were upregulated in arginine relative to no arginine. In biofilms, 922 loci were downregulated and 963 were upregulated in arginine relative to no arginine. Of the loci upregulated in biofilms independent of arginine, most are in the *ebp* operon encoding production of pili subunits. 24 loci were downregulated in biofilms relative to planktonic culture independent of arginine treatment. Of these, 21 are part of the cryptic phage02 operon spanning OG1RF_11046-11061 (34). Many of these proteins were previously identified in membrane vesicles derived from OG1RF (35), but the role of phage02 in biofilm formation is unknown. Of the arginine-responsive loci, 842 were upregulated and 852 were downregulated in both planktonic and biofilm cultures relative to the untreated controls. This suggests that a majority of transcriptional changes that occur during growth in arginine happen independent of growth state (biofilm or planktonic). Based on this, and because we saw significant changes in aggregation during planktonic culture, additional analysis of gene expression changes was done using the planktonic RNAseq dataset.

**Figure 3.**
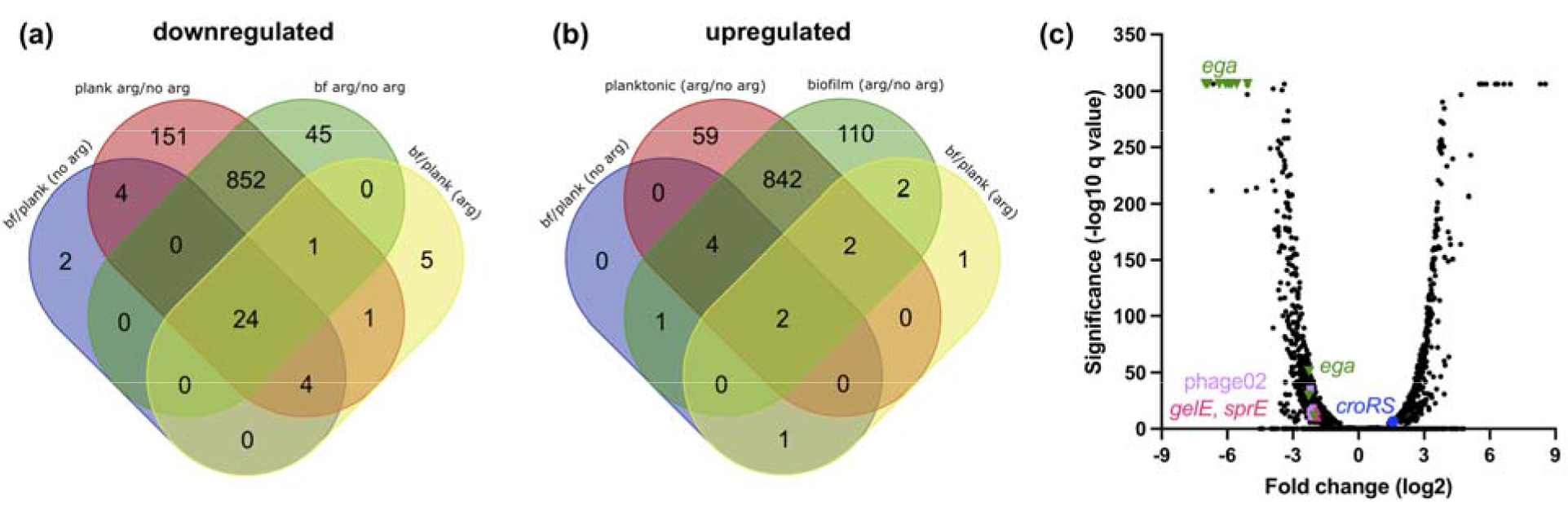
Differential gene expression in OG1RF after growth in arginine. Venn diagrams show **(a)** downregulated and **(b)** upregulated genes after planktonic or biofilm growth with or without arginine. **(c)** Volcano plot highlighting genes differentially expressed during planktonic growth in arginine relative to untreated controls. q values of 0 were set to the lowest q value obtained (E-306) prior to calculating log_10_ for visualization purposes.

The 10 most highly upregulated and downregulated loci during planktonic growth in arginine are shown in **Table 1**. In accordance with previous work (15), expression of *arcA* and *arcB* increased during growth with arginine (**Supplementary Table 1**). The first group of highly downregulated genes, spanning OG1RF_10935-10939, is within the recently named <u>e</u>nteroccocal <u>g</u>lycosyl<u>a</u>sparaginase locus (*ega*) (36). These encode an N4-(β-N- acetylglucosaminyl)-l-asparaginase (EgaG, OG1RF_10935), the EIIB and EIIC subunits of a lactose/cellobiose phosphotransferase system, and a pseudogene that encodes a putative oxidoreductase in other strains (36, 37). OG1RF_10933 and OG1RF_10934, which encode a GntR transcriptional regulator and a dipeptidase and are controlled by a different promoter than OG1RF_10935-10939 (36), were also downregulated, albeit to a lesser extent. The second set of highly downregulated genes includes OG1RF_11941 (encoding glutamate synthase) and OG1RF_11942 (encoding a ferredoxin-NADP+ reductase subunit), which are in an operon (OG1RF_11941-11962) that is involved in selenium and molybdenum metabolism (38) (**Supplementary Table 1**). These genes are not commonly found outside of *E. faecalis* and have been implicated in redox control (3). These gene groups suggest that *E. faecalis* may downregulate metabolic pathways involved in utilizing other nutrients such as sugars or amino acids during growth in arginine.

**Table 1.** Genes over- and underexpressed in OG1RF after planktonic growth in arginine. The 10 most highly over- and underrepresented transcripts (statistically significant with q < 0.05) are shown. Log2FC (log2 fold change) values were calculated based on growth in arginine compared to untreated conditions.

| **<u>Underexpressed in arginine</u>** |  |  |
| --- | --- | --- |
| **<u>Locus tag</u>** | **<u>Gene product</u>** | **<u>Log2FC</u>** |
| Intergenic_988 | Between OG1RF_10936 (*celA4*) and OG1RF_10937 (*celB2*) | -6.99 |
| OG1RF_10936<br>(*celA4*) | PTS system transporter subunit I | -6.89 |
| OG1RF_11942 | Ferredoxin--NADP(+) reductase subunit alpha | -6.70 |
| OG1RF_10939 | Hypothetical protein | -6.64 |
| OG1RF_10937<br>(*celB2*) | PTS family oligomeric beta-glucoside porter component<br>IIC | -6.36 |
| Intergenic_1674 | Between OG1RF_11619 and OG1RF_11620 | -6.25 |
| Intergenic_987 | In between OG1RF_10935 (*ansB*) and OG1RF_10936 (*celA4*) | -6.07 |
| OG1RF_10938 | Hypothetical protein | -5.93 |
| OG1RF_11941 (*gltA*) | Glutamate synthase | -5.82 |
| Intergenic_989 | Between OG1RF_10937 (*celB2*) and OG1RF_10938 | -5.81 |
| **<u>Overexpressed in arginine</u>** |  |  |
| **<u>Locus tag</u>** | **<u>Gene product</u>** | **<u>Log2FC</u>** |
| Intergenic_446 | Between OG1RF_10416 and OG1RF_10417 (*pbp2A*) | 5.52 |
| Intergenic_1362 | Between OG1RF_11307 and OG1RF_11308 | 5.54 |
| OG1RF_10759 | Hypothetical protein | 5.66 |
| OG1RF_11308 | N-acetyltransferase | 5.83 |
| OG1RF_12331 | Lipase | 6.25 |
| OG1RF_11689 | Hypothetical protein | 6.36 |
| OG1RF_10757 (*ppdK*) | Pyruvate phosphate dikinase | 6.64 |
| Intergenic_1750 | Between OG1RF_nc10028 (*glmS*, glucosamine-6-phosphate activated ribozyme), and OG1RF_11693 | 6.93 |
| OG1RF_10535 | Hypothetical protein | 8.30 |
| OG1RF_10328 | Peptidoglycan binding protein | 8.54 |

Unlike the downregulated genes, the genes most highly expressed during growth in arginine were not in operons or clusters in the genome (**Table 1**). The most overexpressed gene, OG1RF_10328, encodes a putative peptidoglycan binding protein. OG1RF_10328 has predicted LysM and DUF4106 domains and is annotated as a peptidoglycan-binding protein (GenBank AEA93015.1). Bacterial LysM-domain proteins, such as a major *E. faecalis* autolysin involved in cell division (AtlA), typically bind N-acetylglucosamine in peptidoglycan (39–41).

OG1RF_10328 was also upregulated in strain V583 (locus tag EF0443) after treatment with antibiotics targeting the cell envelope (42, 43) and is part of the CroRS regulon, which responds to cell wall stress (44, 45). In addition to OG1RF_10328, OG1RF_10535 and OG1RF_12331 were upregulated after exposure to antibiotics that target the cell envelope (42) and in arginine- grown cells. OG1RF_10759 (EF1026) has similarity to a gene product involved in carbon catabolite repression in *Bacillus* (46). Additional genes upregulated during growth in arginine include many involved in virulence and biofilm formation, including the enterococcal polysaccharide biosynthesis operon (*epa*, OG1RF_11706-11738) and *ebpABC* (OG1RF_10869- 10871), which encode the Ebp pili subunits. Curiously, the quorum sensing gene *fsrA* was upregulated, but the Fsr-regulated proteinases GelE and SprE were downregulated. This suggests there may be decoupling of quorum sensing and GelE/SprE expression during growth in arginine. To validate these findings, we used an agar plate-based assay to evaluate gelatinase production of OG1RF grown on CBM agar or CBM supplemented with 25 mM arginine or 25 mM glucose. Colonies grown on plates supplemented with arginine produced gelatinase zones with smaller diameters than cells grown on unsupplemented plates or plates with glucose (**Supplementary** Figure 2). This reduced GelE activity supports the changes in *gelE* gene expression observed using RNAseq.

### Comparison of *E. faecalis* growth in arginine and alkaline stress

Ammonia generation via the ADI pathway during growth in arginine could put cells under alkaline stress, and previous work found that growth of *E. faecalis* at pH 10 resulted in biofilms with less surface coverage relative to unadjusted medium (15, 47). We measured the pH of our *E. faecalis* cultures grown in untreated CBM or medium supplemented with arginine, glycine, and glucose. After 24 hr, the pH of cultures grown in 25 or 50 mM arginine increased from pH 7 to approximately pH 9 (**Supplementary** Figure 3). Although *E. faecalis* is highly tolerant to pH stress (48), we were curious whether any differential gene expression in our RNAseq could be attributed to increased pH. To determine this, we compared our data to previously published studies on gene expression and adaptation to growth at high pH. First, we examined a study of the transcriptional response of *E. faecalis* ATCC 33186 to growth at pH 10 (47). Of the 613 genes differentially expressed at pH 10, we identified 253 as also differentially regulated in our arginine RNAseq (**Supplementary Table 2**). However, only 33 genes had the same pattern of gene expression (up or downregulated in both studies). The 17 commonly upregulated genes included 7 encoding hypothetical proteins and 4 encoding ribosomal proteins. The 16 commonly downregulated genes included those involved in pyrimidine metabolism (OG1RF_11425-11429) and a putative iron-siderophore ABC transporter (OG1RF_12351- 12354). Next, we compared our results to studies showing that growth at high pH increased expression of 37 *E. faecalis* proteins, including the heat-shock proteins DnaK and GroEL, and genes including *ace*, *fsrB*, and *gelE* (49, 50). In our work, expression of the heat-shock operon (OG1RF_11076-11080, *hrcA*-*dnaJ*), *ace* (OG1RF_10878), and *fsrB* (OG1RF_11528) were not significantly changed after growth in arginine, and expression of *gelE* (OG1RF_11526) was lower (**Supplementary Table 1**). Finally, we compared our arginine RNAseq to a study that identified mutations in base-adapted *E. faecalis* OG1RF (51). We found little correlation between these mutations and the gene expression pattern in RNAseq. Multiple high pH-evolved clones had mutations in genes encoding the Opp peptide transport system, the Pst phosphate transport system, OG1RF_11160 (encoding a putative thioesterase), and CcpA (51). In our work, *oppABCDF* and OG1RF_11160 were upregulated, *ccpA* was downregulated, and there was no change in expression of the *pst* operon. Together, this suggests that some changes in gene expression after growth in arginine may be due to alkaline stress, but the majority of changes we measured using RNAseq did not correlate with known patterns of gene or protein expression in high pH. As such, these changes in gene expression may be specifically linked to arginine catabolism and not simply a change in culture pH.

### Arginine affects antibiotic susceptibility and cell envelope permeability of *E. faecalis*

In *Streptococcus pyogenes*, the ADI pathway affects biofilm-associated antibiotic resistance (24). Therefore, we searched our RNAseq data to see whether any differentially expressed genes were linked to antibiotic resistance in *E. faecalis*. The *liaFSR* genes associated with daptomycin resistance (52, 53) were significantly downregulated after growth in arginine, but many genes involved in resistance to antibiotics that target the cell envelope, such as those encoding penicillin-binding proteins, were upregulated (**Table 2**). Expression of *croRS*, which mediates cephalosporin resistance (44, 54), was moderately upregulated, but expression of *ireK* was not significantly changed.

**Table 2.** Antibiotic resistance genes differentially expressed in *E. faecalis* OG1RF after growth in arginine. For genes where log2FC values are shown, q < 0.05.

| Locus tag (gene name) | Log2FC |
| --- | --- |
| OG1RF_10417 (*pbp2A*) | 3.91 |
| OG1RF_10724 (*pbpC*) | 3.83 |
| OG1RF_10925 (*pbp1A*) | 3.21 |
| OG1RF_11450 (*pbp1B*) | 2.39 |
| OG1RF_11907 (*pbp4(5)*) | 1.28 |
| OG1RF_12158 (*penA/pbpA(2B)*) | 1.98 |
| OG1RF_12165 (*aadK*) | 3.79 |
| OG1RF_12193 | 1.33 |
| OG1RF_12211 (*liaR*) | -1.89 |
| OG1RF_12212 (*liaS*) | -1.47 |
| OG1RF_12213 (*liaF*) | -1.87 |
| OG1RF_12535 (*croR*) | 1.54 |
| OG1RF_12536 (*croS*) | 1.48 |

Based on this, we hypothesized that antibiotic susceptibility would be altered in arginine- grown *E. faecalis* relative to cells grown in untreated medium and tested this using disk diffusion assays (**Figure 4A**). OG1RF was grown on agar plates supplemented with arginine or glucose in the presence of a panel of antibiotics (ampicillin, cephalothin, ceftriaxone, gentamicin, linezolid, penicillin, minocycline, and vancomycin). Relative to growth on glucose or untreated CBM, we observed smaller zones of clearance (indicating less susceptibility) for ampicillin and ceftriaxone, although only the difference in ampicillin was statistically significant (**Figure 4B**). This is consistent with the increased expression of *croRS* in the RNAseq, as deletion of *croRS* in the strain JH2-2 led to greater susceptibility to ampicillin and ceftriaxone (54). We did not detect a change in susceptibility to minocycline and linezolid, which target the ribosome. These results suggest that susceptibility to antibiotics targeting the cell envelope may be altered in arginine-rich environments such as the oral cavity or gastrointestinal tract.

**Figure 4.**
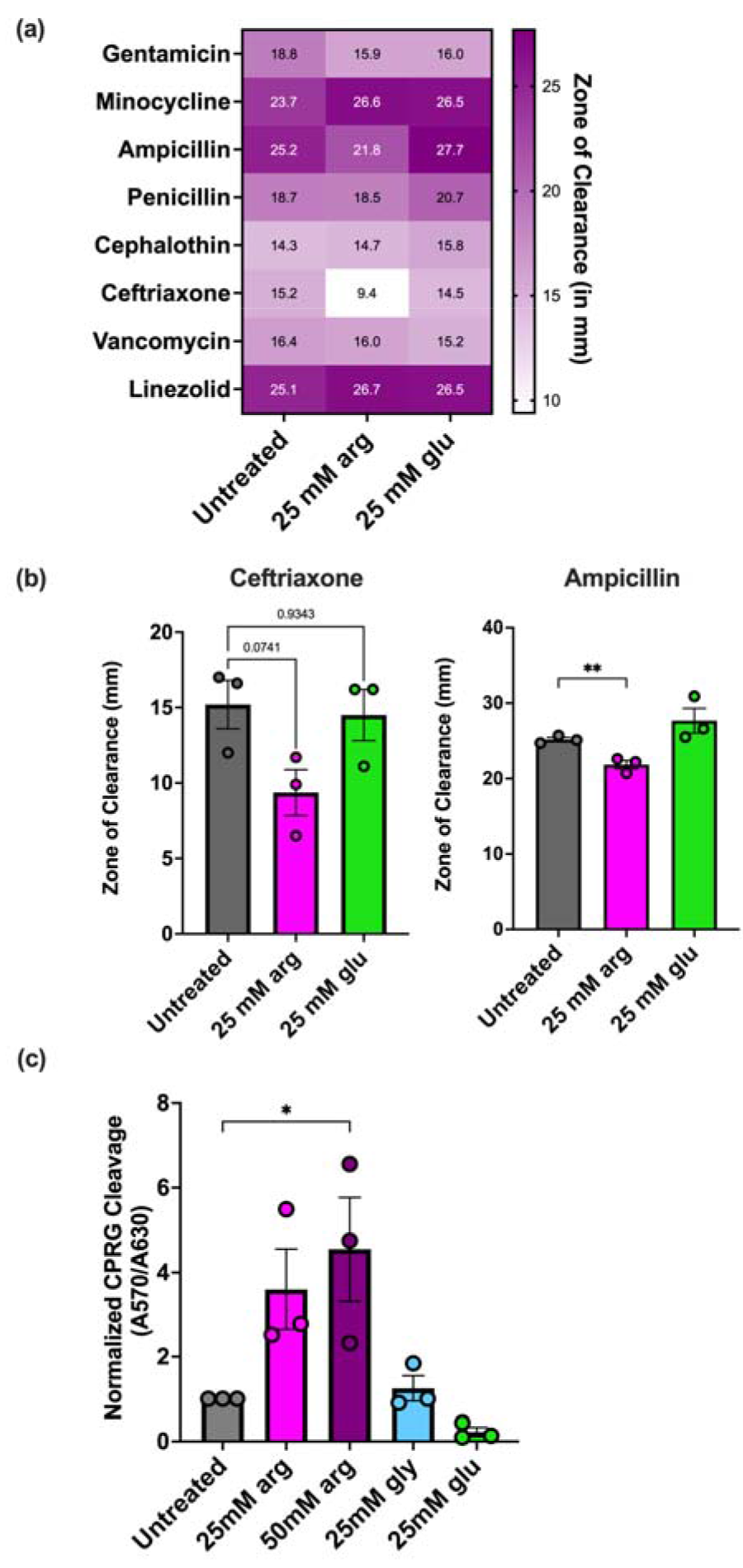
Growth with arginine affects cell envelope permeability and antibiotic resistance in *E*.*faecalis*. **(a)** Heatmap of zones of inhibition (measured in mm) of antibiotic disk diffusion assay. Cells were grown on CBM agar plates supplemented with arginine or glucose (25 mM). Disks measured 9 mm in diameter. Data represents the average of three independent biological replicates. **(b)** Individual values of disk diffusion zones for ceftriaxone and ampicillin. **(c)** Generation of the CPRG cleavage product (A570) relative to growth (A600) normalized to untreated cultures. For panels (b) and (c), each data point represents an independent biological replicate. Statistical significance was determined using one-way ANOVA with Dunnett’s test for multiple comparisons. *p<0.05, **p<0.01.

Given the changes in gene expression and antibiotic susceptibility after growth in arginine, we next asked whether integrity of the cell envelope was impacted in arginine-grown cells. To test this, we carried out a cell envelope permeability assay in which the cell-impermeable LacZ substrate chlorophenol red-β-D- galactopyranoside (CPRG) is added to cultures of cells constitutively expressing *lacZ* from a plasmid. Cleavage of CPRG, which produces a color change and can be measured by absorbance at 570 nm, can occur when the cell envelope is compromised (55, 56). We grew OG1RF in arginine, glycine, glucose, or untreated medium and quantified CPRG cleavage after 24 hr growth (**Figure 4C**). In 50 mM arginine, there was a significant increase in CPRG cleavage relative to the untreated control. There was no statistically significant change with cultures grown in glycine or glucose. Together, this data demonstrates that growth in arginine changes antibiotic susceptibility and cell envelope permeability of *E. faecalis*.

## Discussion

Arginine metabolism by *E. faecalis* affects polymicrobial interactions and promotes virulence of other pathogens (10, 13). Our work demonstrates that arginine also induces widespread changes in gene expression, biofilm formation, and antibiotic resistance in *E. faecalis*. Similar to other Gram-positive bacteria, *E. faecalis* biofilm formation decreased in the presence of arginine, which could have implications for how *E. faecalis* forms biofilms in arginine-rich environments like the oral cavity or GIT. However, pre-formed *E. faecalis* biofilms were not disrupted by the same concentrations of arginine that reduced biofilm growth. Additionally, high concentrations of arginine (250-500 mM) that were previously shown to destabilize *S. mutans* biofilms (29) did not disrupt pre-grown *E. faecalis* biofilms as measured by safranin staining. We also observed differences in gene expression of arginine-grown *E. faecalis* compared to other oral bacteria. In *S. mutans*, expression of the *dnaK* stress response operon was upregulated after growth in 1.5% arginine (28). However, in our RNAseq, expression of the *dnaJK* stress response genes were not significantly changed during growth in arginine. This could potentially be relevant for oral health as arginine is used in dentistry to control plaque and prevent caries (cavities) (18). *E. faecalis* is not considered part of the normal oral microbiome but is a leading cause of infected root canals (57, 58), so it is interesting to consider whether the relative abundance of *E. faecalis* in the oral cavity would be affected by arginine treatment in healthy patients or those with active root canal infections. Given that we measured a decrease in antibiotic susceptibility of *E. faecalis* during growth in arginine, it is also interesting to consider how the use of arginine in the oral cavity could affect virulence of *E. faecalis*. We observed arginine-related *E. faecalis* phenotypes at relatively low concentrations (25 and 50 mM) compared to other studies examining the impact of arginine treatment on the oral microbiome, co-culture biofilm structure and composition, exopolysaccharide, and gene expression (21, 29). Therefore, it is possible that *E. faecalis* may respond to arginine at sub-clinical levels relative to what is used in dentistry applications. However, the broader impact of how this affects *E. faecalis* biofilms and antibiotic resistance in the oral cavity or other body sites is yet undetermined.

Arginine metabolism can confer benefits to bacteria during adaptation to a mammalian niche. We found some overlap in arginine-induced gene expression changes and other studies on *E. faecalis* growth in alkaline stress (47, 49, 50), and previous work showed that alkaline-adapted *E. faecalis* acquired cross-resistance to bile salts (49). Therefore, arginine exposure in the oral cavity or GIT (and the resulting changes in pH and gene expression) could prime *E. faecalis* for survival against bile salts and other stressors in the GIT. ADI pathway-derived ammonia was hypothesized to promote survival of *Staphylococcus epidermidis* during biofilm growth, as expression of the ADI pathway was upregulated in biofilms (59). Additionally, arginine metabolism through an ADI pathway encoded on a mobile element in *S. aureus* USA300 was important for survival in skin-mimicking acidic conditions (60). ADI pathways were identified in mammalian-associated Saccharibacteria but not those isolated from environmental habitats, and the presence of arginine maintained infectivity of *Nanosynbacter lyticus* strain TM7x in conditions that mimicked the oral cavity (61). However, the presence of arginine promoted cell membrane integrity in TM7x whereas we demonstrated here that growth in arginine disrupts cell envelope integrity in *E. faecalis.* Together, this suggests that while the ADI pathway may promote survival of diverse bacteria *in vivo*, the mechanisms by which this occurs can differ across species.

The ADI pathway has previously been linked to antibiotic tolerance during oxidative stress and biofilm growth in *E. faecalis* and other bacteria. Upregulation of the ADI operon was associated with restored vancomycin tolerance in *E. faecalis* mutants lacking *sodA* (encoding superoxide dismutase), and disruption of the ADI pathway in a *sodA* mutant background led to increased killing by vancomycin relative to the *sodA* mutant alone (62). In *S. pyogenes*, the ADI pathway is important for biofilm-associated antibiotic resistance *in vitro* and in an *in vivo* animal model (24). Here, we found changes in the expression of multiple genes linked to *E. faecalis* antibiotic tolerance and found that *E. faecalis* grown with arginine was less susceptible than control cells to antibiotics targeting the cell envelope. Based on this, it is interesting to speculate that arginine metabolism in various body sites may contribute to tolerance of *E. faecalis* to antibiotics, and whether this could contribute to persistence or overgrowth during antibiotic treatment.

Previous work with *S. gordonii* showed that treatment with exogenous arginine, but not glycine, weakened biofilms and led to increased removal by shear stress (22). Although that work used higher concentrations (4% arginine and 0.23 M glycine) than our studies, we also observed no change in aggregation or biofilm formation in the presence of glycine. The exception to this was an apparent increase in biofilm index when pre-formed early biofilms (6 hr) were treated with glycine. We speculate that this could be due to interactions with glycine and components of the early biofilm matrix, given that this increase was not observed with 24 hr biofilms, and previous work indicates there are temporal differences in genetic determinants of biofilm formation (17, 32). Additional outstanding questions are the mechanisms by which growth in arginine leads to changes in biofilm morphology. *S. mutans* biofilms grown with 1.5% arginine had reduced levels of exopolysaccharides and multi-cellular clusters or microcolonies (21). Conversely, we observed an increase in *E. faecalis* aggregation during planktonic and biofilm growth in arginine, although morphology changes are also evident in glucose-grown biofilms compared to untreated media. In our RNAseq, we found increased expression of genes in the Epa operon, which encodes for biosynthesis of the cell wall-associated enterococcal polysaccharide antigen (63, 64). Epa modifications can alter *E. faecalis* biofilm morphology, leading to an increase in multi-cellular clusters or microcolonies instead of the flat monolayer biofilms typically formed *in vitro* by *E. faecalis* OG1RF (31, 33). However, more work is needed to understand what specifically drives arginine-induced biofilm morphology rearrangement and how that might translate to biofilm remodeling *in vivo*.

Our work shows that *E. faecalis* growth and biofilm formation is altered in the presence of arginine. Importantly, we also identified species-specific differences in expression of stress response genes, biofilm destabilization, and biofilm architecture relative to arginine-induced changes in Gram-positive bacteria. Together, this work provides a better understanding of how *E. faecalis* responds to growth in arginine independent of cross-feeding that promotes virulence of other pathogens and provides a platform from which the impact of *E. faecalis* gene expression changes on polymicrobial interactions and infections can be studied in addition to phenotypes mediated via arginine metabolism through the ADI pathway.

## Materials and Methods

### Bacterial strains, growth conditions, and reagents

All strains used are listed in **Supplementary Table 3**. Bacterial stocks were maintained at -80 °C in 25% glycerol (w/v). Strains were routinely grown in complex basal medium (CBM) (14) or brain-heart infusion broth (BHI, Difco). Media was supplemented with 1% agar for solid plates. Unless otherwise noted, strains were cultured under static conditions. Where indicated, cultures were supplemented with arginine, glucose, and glycine (Sigma) from 1 M stock solutions that were prepared in water and filter sterilized. Antibiotic disks (BD BBL) were purchased from Fisher Scientific.

### RNA sequencing and analysis

OG1RF was diluted to OD600 = 0.02 and grown in CBM with or without 25 mM arginine for 6 hr. Two independent biological replicates were used for each condition. Planktonic cells (∼2x10^9^ CFU) were mixed with a 2:1 volume of RNAprotect (Qiagen), pelleted, and stored at -80 °C. Biofilms were grown on Aclar coupons, which were rinsed in phosphate-buffered saline with 10 mM potassium at pH 7.4 and scraped with a razorblade to remove cells. Cells were pelleted and stored at -80 °C. Pellets were resuspended in 200 μL TE buffer (10 mM Tris pH 8.0, 1 mM EDTA) supplemented with 30 mg/mL lysozyme and 500 U/mL mutanolysin and incubated at 37 °C for 10 min, after which RNA was extracted using a Qiagen RNeasy kit following manufacturer’s instructions. Lysozyme and mutanolysin were purchased from Sigma. Residual genomic DNA was removed using Turbo DNase (Ambion) following the manufacturer’s protocol for two DNase treatments (rigorous method).

Complete removal of DNA was confirmed using PCR with oligos JD460s and JD461as (**Supplementary Table 3**). rRNA was depleted using MICROBExpress (Ambion). RNA was sent to the University of Minnesota Genomics Center for TruSeq stranded library preparation and sequencing on a single lane of an Illumina NextSeq 550 in mid-output mode (75-bp paired-end reads). Sequencing quality was evaluated using FastQC (65). Reads were trimmed with Trimmomatic (66) and imported into Rockhopper for analysis using standard settings (67, 68). Reads were aligned against a custom OG1RF reference genome in which intergenic regions are annotated. Fold changes (log2FC) were calculated from Rockhopper expression values, and q-values ≤ 0.05 were considered statistically significant.

### Growth measurements

For growth curves, OG1RF cultures were grown overnight in 3 mL of CBM. Cells were diluted 1:100 in a 96-well plate (Corning 3595) using untreated CBM or CBM supplemented with arginine (25 or 50 mM), glucose (25 mM), glycine (25 or 50 mM), or a vehicle control for a final volume of 200 μL. Three independent biological replicates (each with 3 technical replicates) were performed. Plates were sealed (Microseal B plate seals, Biorad) and incubated in a Biotek Epoch 2 microplate reader without shaking at 37 °C for 15 hr. The A_600_ was measured every 30 minutes. Technical replicates were averaged, and the average of all biological replicates was plotted. For quantification of colony forming units, OG1RF cultures were grown in glass culture tubes in 3 mL CBM supplemented with arginine, glycine, and glucose (at 25 mM and 50 mM). After 24 hr, cultures were diluted (10-fold serial dilutions) and spotted onto agar plates. CFU/mL was quantified after 24 hr incubation.

### Biofilm assays

Cultures were grown overnight at 37 °C in 3 mL of CBM. Cells were diluted 1:50 in a 96-well plate (Corning 3595) in 200 μL of CBM supplemented with arginine or glycine (25 or 50 mM). Three independent biological replicates were done for each condition. Plates were incubated in a humidified plastic container at 37 °C for 6 or 24 hr. Growth of the cells was quantified by measuring the A_600_ using a Biotek Epoch 2 microplate reader. The planktonic cells and media were removed by washing 3x with de-ionized water. After drying, the plates were than stained with 0.1% w/v safranin (Sigma) for 20 minutes, at which point the plates were washed again with de-ionized water. Stained biofilm material was quantified by measuring A_450_. The biofilm index was then calculated as A_450_/A_600_ as a representation of biofilm biomass relative to planktonic cell growth.

### Fluorescence microscopy of biofilms

OG1RF cultures were grown overnight in 3 mL of CBM at 37 °C and diluted 1:50 into an optically clear 96-well plate (Nunc MicroWell tissue culture-treated optical bottom plates). Each well contained 200 μL of CBM with or without arginine, (25 mM and 50 mM), glucose (25 mM and 50 mM), or glycine (25 mM and 50 mM). Plates were grown in a humidified chamber at 37 °C for 24 hours. Plates were then washed 1x with phosphate buffered saline (PBS, pH 7.4) and 200 μL of formalin (10% w/v) was added at 4 °C

for 16 hr to fix the cells. Formalin was removed with a multichannel pipette, and the plate then was washed once with PBS. 200 μL of Hoechst 33342 (Thermo Fisher Scientific) dye was added (final concentration 5 μg/mL) and incubated on the lab bench for 30 minutes prior to imaging. The plate was then imaged using a Keyence BZ-X810 microscope with a Chroma DAPI filter (AT350/50x) at 20x magnification. At least three images were acquired for each independent biological replicate. Images were processed using the rolling ball background subtraction feature in Fiji and cropped to 500 x 500 pixels.

### Auto-aggregation assay

Strains were diluted 1:100 in 2 mL of CBM supplemented with arginine (25 or 50 mM), glycine (25 mM), or glucose (25 mM). Cultures were grown for 24 hours at 37 °C at which point 100 μL was removed from the top of the culture, mixed with 900 μL medium, and used to measure A_600_. Cultures were vortexed, and another A_600_ reading was taken for the mixed culture. To quantify aggregation, the A_600_ of the unmixed culture was divided by the A_600_ culture after mixing and multiplied by 100. Treatment conditions were normalized to untreated cultures.

### Cell envelope permeability assay

OG1RF pCJK205 was grown overnight in 3 mL of CBM with 10 μg/mL of erythromycin (Erm) at 37 °C. The culture was then diluted to an A_600_ of 0.05 in 5 mL of fresh CBM, 10 μg/mL Erm, and 20 μg/mL of chlorophenol red-β-D-galactopyranoside (CPRG) supplemented with arginine (25 or 50 mM), glycine (25 mM), or glucose (25 mM).

Cultures were incubated for 24 hours at 37 °C, at which time the A_630_ was measured. 750 μL of culture was pelleted, and the A_570_ of the supernatant was measured. To quantify CPRG cleavage relative to growth, the A_570_ value was divided by the A_600_. Results were normalized to untreated OG1RF.

### Antibiotic disc diffusion assay

CBM agar plates were supplemented with 25 mM arginine, 25 mM glucose, or left untreated. Cultures of OG1RF were grown overnight in CBM at 37 °C, and 200 μL was spread onto each plate. Plates were incubated at room temperature for 15 min after which antibiotic discs (gentamicin, ampicillin, penicillin, minocycline, vancomycin, cephalothin, ceftriaxone, and linezolid) were placed on the plate surface. Plates were then incubated for 24 hours at 37 °C, after which plates were imaged with a Cell Biosciences FluorChem FC3 plate

reader. The zone of clearance for each antibiotic was measured in millimeters using Fiji (69).

### Gelatinase activity assay

Overnight cultures of OG1RF ere grown in 3 mL CBM at 37 °C. 5 μL was spotted on CBM agar plates supplemented with 3% w/v gelatin and 25 mM arginine or glucose. Plates were incubated at 37 °C for 24 hr, after which they were transferred to 4 °C for 24 hr prior to imaging. Images were taken using a Cell Biosciences FluorChem FC3 images, and the diameter of each gelatinase halo was measured in millimeters using Fiji (69).

### Statistical analysis

Statistical analysis was performed using GraphPad Prism (version 10.1.1). Statistical tests and the number of replicates represented by each data point are described in figure legends.

### Data availability

Raw sequencing files and processed data have been deposited in the NCBI GEO database with accession number GSE268264.

## Acknowledgements

We thank the Dunny and Willett labs for helpful input on the manuscript. This work was supported by National Institutes of Health grant R01AI122742 to GMD and start-up funds from the University of Minnesota to JLEW. The content is solely the responsibility of the authors and does not necessarily represent the official views of the National Institutes of Health. This work was also supported by the resources and staff at the University of Minnesota Genomics Center (RRID: SCR_012413).

---

**Supplementary Figure 1.**
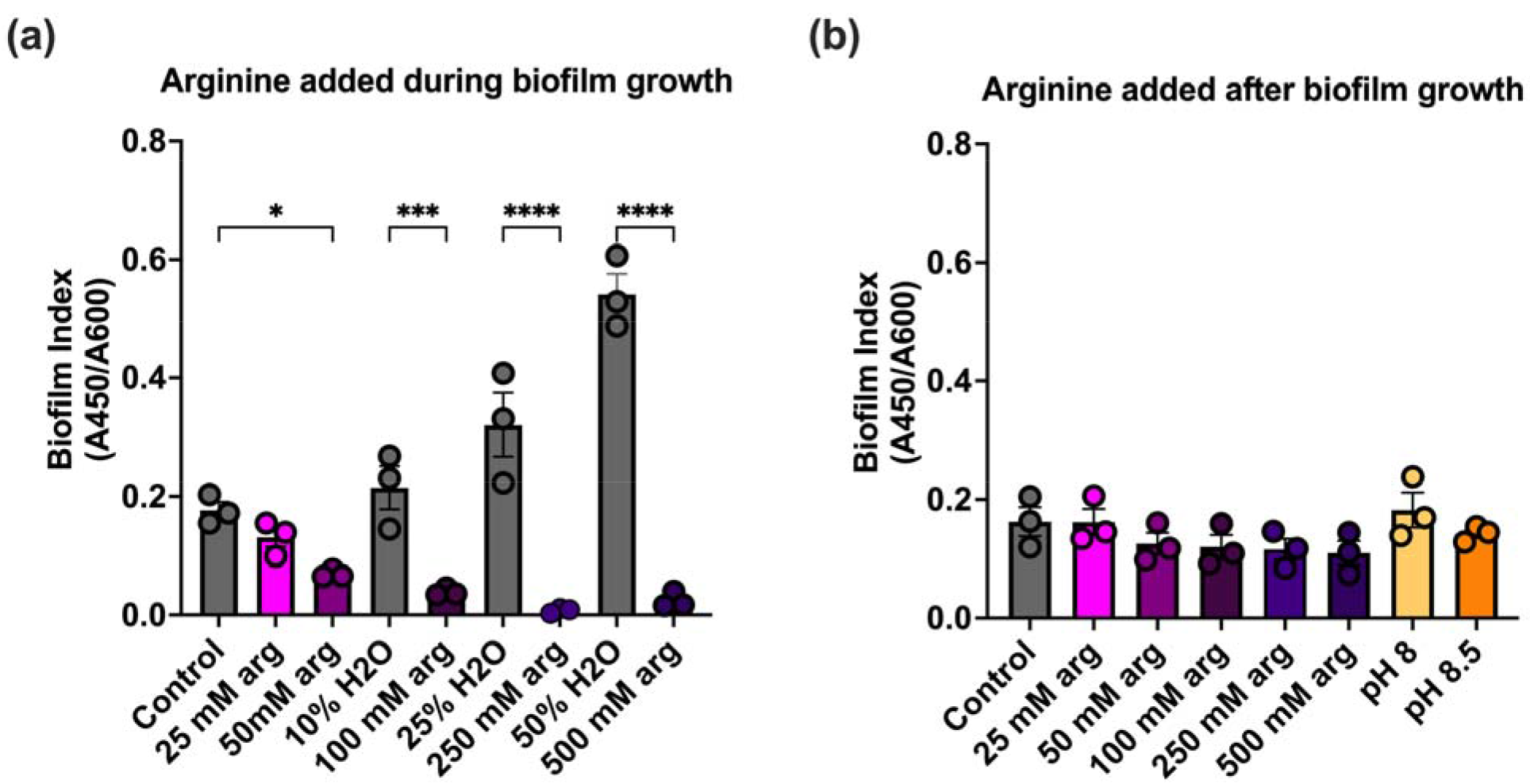
*E. faecalis* biofilm formation in high concentrations of arginine. **(a)** *E. faecalis* OG1RF was grown for 24 hr with arginine added at the indicated concentrations. Due to the increasing volume of arginine in samples treated during growth, each arginine-treated sample is shown next to a volumetric control. **(b)** Biofilms were grown in the absence of arginine and washed, after which arginine, vehicle control, or Tris-HCl buffer at the indicated concentration were added for 20 min. For both panels, each data point represents an individual biological replicate. Statistical significance was evaluated using one-way ANOVA with Sidak’s test (in panel **(a)**) or Dunnett’s test (in panel **(b)**) for multiple comparisons. Comparisons that were not statistically significant are not shown. *p<0.05,**p<0.01, ***p<0.001, ****p<0.0001.

**Supplementary Figure 2.**
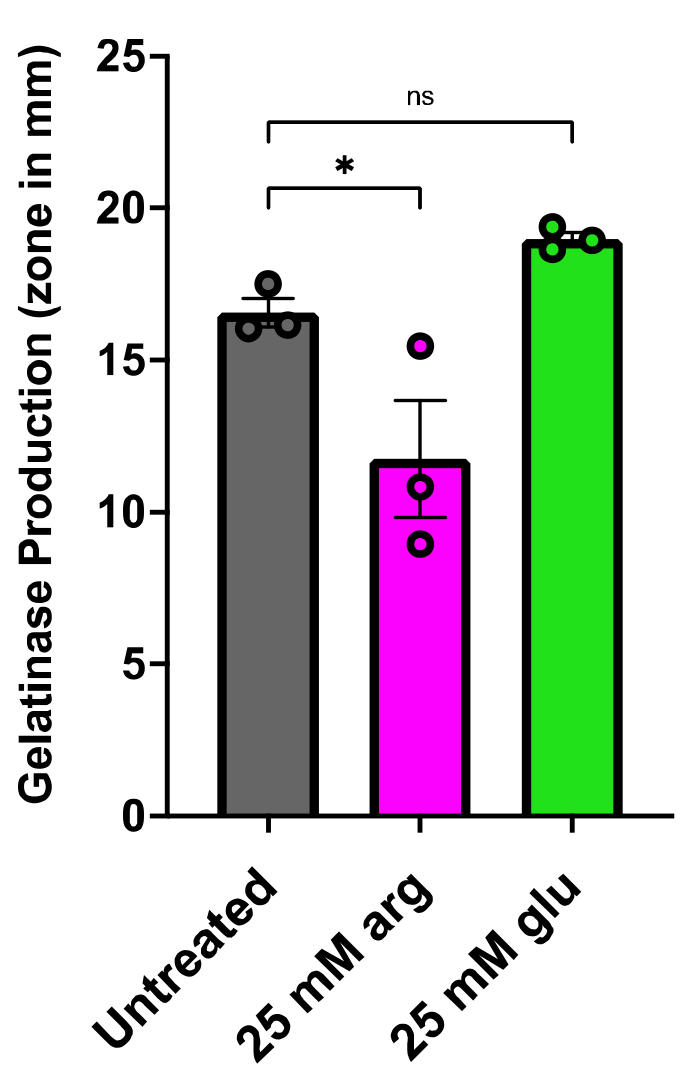
Gelatinase activity of *E. faecalis* OG1RF grown with or without arginine. *E. faecalis* OG1RF cultures were spotted onto agar plates supplemented with 3% gelatin and 25 mM arginine or glucose. Hazy zones indicative of gelatinase activity were measured in mm after 24 hr growth. Each data point represents an individual biological replicate. Statistical significance was determined by one-way ANOVA with Dunnett’s test for multiple comparisons. *p<0.05.

**Supplementary Figure 3.**
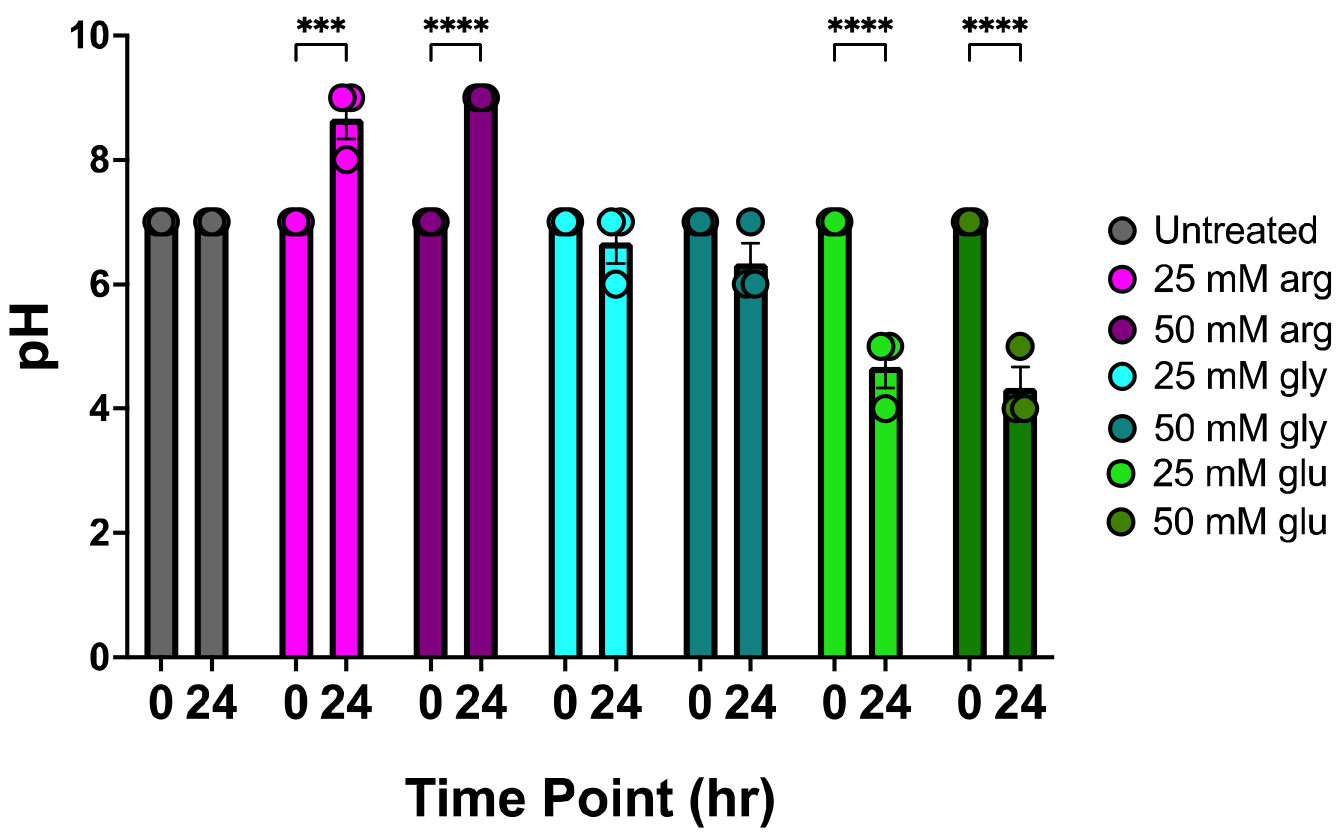
pH of *E. faecalis* cultures. *E. faecalis* OG1RF was cultured in CBM supplemented with the indicated nutrients. pH measurements were taken at 0 hr prior to growth and after 24 hr incubation. Each data point represents an independent biological replicate. Error bars represent standard deviation. Statistical significance between 0 and 24 hr for each condition was assessed with two-way ANOVA and Sidak’s test for multiple comparisons. ***p<0.001, ****p<0.0001

## References

1. Gilmore MS, Lebreton F, van Schaik W. Genomic transition of enterococci from gut commensals to leading causes of multidrug-resistant hospital infection in the antibiotic era. Curr Opin Microbiol. 2013;16(1):10–6.

2. Lebreton F, Manson AL, Saavedra JT, Straub TJ, Earl AM, Gilmore MS. Tracing the Enterococci from Paleozoic Origins to the Hospital. Cell. 2017;169(5):849–61.e13.

3. Gilmore MS, Salamzade R, Selleck E, Bryan N, Mello SS, Manson AL, et al. Genes Contributing to the Unique Biology and Intrinsic Antibiotic Resistance of Enterococcus faecalis. mBio. 2020;11(6).

4. Tan CAZ, Antypas H, Kline KA. Overcoming the challenge of establishing biofilms in vivo: a roadmap for Enterococci. Curr Opin Microbiol. 2020;53:9–18.

5. Goh HMS, Yong MHA, Chong KKL, Kline KA. Model systems for the study of Enterococcal colonization and infection. Virulence. 2017;8(8):1525–62.

6. García-Solache M, Rice LB. The Enterococcus: a Model of Adaptability to Its Environment. Clin Microbiol Rev. 2019;32(2).

7. Tien BYQ, Goh HMS, Chong KKL, Bhaduri-Tagore S, Holec S, Dress R, et al. Enterococcus faecalis Promotes Innate Immune Suppression and Polymicrobial Catheter-Associated Urinary Tract Infection. Infect Immun. 2017;85(12).

8. Tan CAZ, Lam LN, Biukovic G, Soh EY, Toh XW, Lemos JA, et al. Enterococcus faecalis Antagonizes Pseudomonas aeruginosa Growth in Mixed-Species Interactions. J Bacteriol. 2022;204(7):e0061521.

9. Ch’ng JH, Muthu M, Chong KKL, Wong JJ, Tan CAZ, Koh ZJS, et al. Heme cross-feeding can augment Staphylococcus aureus and Enterococcus faecalis dual species biofilms. ISME J. 2022;16(8):2015–26.

10. Smith AB, Jenior ML, Keenan O, Hart JL, Specker J, Abbas A, et al. Enterococci enhance Clostridioides difficile pathogenesis. Nature. 2022;611(7937):780-6.

11. Gaston JR, Andersen MJ, Johnson AO, Bair KL, Sullivan CM, Guterman LB, et al. *Enterococcus faecalis* Polymicrobial Interactions Facilitate Biofilm Formation, Antibiotic Recalcitrance, and Persistent Colonization of the Catheterized Urinary Tract. Pathogens. 2020;9(10).

12. Gaston JR, Johnson AO, Bair KL, White AN, Armbruster CE. Polymicrobial interactions in the urinary tract: is the enemy of my enemy my friend? Infect Immun. 2021.

13. Keogh D, Tay WH, Ho YY, Dale JL, Chen S, Umashankar S, et al. Enterococcal Metabolite Cues Facilitate Interspecies Niche Modulation and Polymicrobial Infection. Cell Host Microbe. 2016;20(4):493–503.

14. Deibel RH. Utilization of arginine as an energy source for the growth of Streptococcus faecalis. J Bacteriol. 1964;87(5):988–92.

15. Barcelona-Andrés B, Marina A, Rubio V. Gene structure, organization, expression, and potential regulatory mechanisms of arginine catabolism in Enterococcus faecalis. J Bacteriol. 2002;184(22):6289–300.

16. Manias DA, Dunny GM. Expression of Adhesive Pili and the Collagen-Binding Adhesin Ace Is Activated by ArgR Family Transcription Factors in Enterococcus faecalis. J Bacteriol. 2018;200(18).

17. Willett JL, Ji M, Dunny GM. Exploiting biofilm phenotypes for functional characterization of hypothetical genes in Enterococcus faecalis. npj Biofilms and Microbiomes volume2019.

18. Nascimento MM. Potential Uses of Arginine in Dentistry. Adv Dent Res. 2018;29(1):98–103.

19. Burne RA, Marquis RE. Alkali production by oral bacteria and protection against dental caries. FEMS Microbiol Lett. 2000;193(1):1–6.

20. Kolderman E, Bettampadi D, Samarian D, Dowd SE, Foxman B, Jakubovics NS, et al. L-arginine destabilizes oral multi-species biofilm communities developed in human saliva. PLoS One. 2015;10(5):e0121835.

21. He J, Hwang G, Liu Y, Gao L, Kilpatrick-Liverman L, Santarpia P, et al. l-Arginine Modifies the Exopolysaccharide Matrix and Thwarts Streptococcus mutans Outgrowth within Mixed-Species Oral Biofilms. J Bacteriol. 2016;198(19):2651–61.

22. Gloag ES, Wozniak DJ, Wolf KL, Masters JG, Daep CA, Stoodley P. Arginine Induced *Streptococcus gordonii* Biofilm Detachment Using a Novel Rotating-Disc Rheometry Method. Front Cell Infect Microbiol. 2021;11:784388.

23. Nascimento MM, Browngardt C, Xiaohui X, Klepac-Ceraj V, Paster BJ, Burne RA. The effect of arginine on oral biofilm communities. Mol Oral Microbiol. 2014;29(1):45–54.

24. Freiberg JA, Le Breton Y, Harro JM, Allison DL, McIver KS, Shirtliff ME. The Arginine Deiminase Pathway Impacts Antibiotic Tolerance during Biofilm-Mediated Streptococcus pyogenes Infections. mBio. 2020;11(4).

25. Manna AC, Leo S, Girel S, González-Ruiz V, Rudaz S, Francois P, et al. Teg58, a small regulatory RNA, is involved in regulating arginine biosynthesis and biofilm formation in Staphylococcus aureus. Sci Rep. 2022;12(1):14963.

26. Dunny G, Funk C, Adsit J. Direct stimulation of the transfer of antibiotic resistance by sex pheromones in Streptococcus faecalis. Plasmid. 1981;6(3):270–8.

27. Jakubovics NS, Robinson JC, Samarian DS, Kolderman E, Yassin SA, Bettampadi D, et al. Critical roles of arginine in growth and biofilm development by Streptococcus gordonii. Mol Microbiol. 2015;97(2):281–300.

28. Chakraborty B, Burne RA. Effects of Arginine on Streptococcus mutans Growth, Virulence Gene Expression, and Stress Tolerance. Appl Environ Microbiol. 2017;83(15).

29. Zheng X, He J, Wang L, Zhou S, Peng X, Huang S, et al. Ecological Effect of Arginine on Oral Microbiota. Sci Rep. 2017;7(1):7206.

30. Leuck AM, Johnson JR, Dunny GM. A widely used in vitro biofilm assay has questionable clinical significance for enterococcal endocarditis. PLoS One. 2014;9(9):e107282.

31. Korir ML, Dale JL, Dunny GM. Role of e*paQ*, a Previously Uncharacterized Enterococcus faecalis Gene, in Biofilm Development and Antimicrobial Resistance. J Bacteriol. 2019;201(18).

32. Willett JLE, Dale JL, Kwiatkowski LM, Powers JL, Korir ML, Kohli R, et al. Comparative Biofilm Assays Using Enterococcus faecalis OG1RF Identify New Determinants of Biofilm Formation. mBio. 2021;12(3):e0101121.

33. Dale JL, Nilson JL, Barnes AMT, Dunny GM. Restructuring of Enterococcus faecalis biofilm architecture in response to antibiotic-induced stress. NPJ Biofilms Microbiomes. 2017;3:15.

34. McBride SM, Fischetti VA, Leblanc DJ, Moellering RC, Gilmore MS. Genetic diversity among Enterococcus faecalis. PLoS One. 2007;2(7):e582.

35. Afonina I, Tien B, Nair Z, Matysik A, Lam LN, Veleba M, et al. The composition and function of *Enterococcus faecalis* membrane vesicles. Microlife. 2021;2:uqab002.

36. Combret V, Rincé I, Budin-Verneuil A, Muller C, Deutscher J, Hartke A, et al. Utilization of glycoprotein-derived N-acetylglucosamine-L-asparagine during Enterococcus faecalis infection depends on catabolic and transport enzymes of the glycosylasparaginase locus. Res Microbiol. 2023:104169.

37. Wagner M, Sonntag D, Grimm R, Pich A, Eckerskorn C, Söhling B, et al. Substrate-specific selenoprotein B of glycine reductase from Eubacterium acidaminophilum. Biochemical and molecular analysis. Eur J Biochem. 1999;260(1):38–49.

38. Zhang Y, Turanov AA, Hatfield DL, Gladyshev VN. In silico identification of genes involved in selenium metabolism: evidence for a third selenium utilization trait. BMC Genomics. 2008;9:251.

39. Buist G, Steen A, Kok J, Kuipers OP. LysM, a widely distributed protein motif for binding to (peptido)glycans. Mol Microbiol. 2008;68(4):838–47.

40. Eckert C, Lecerf M, Dubost L, Arthur M, Mesnage S. Functional analysis of AtlA, the major N-acetylglucosaminidase of Enterococcus faecalis. J Bacteriol. 2006;188(24):8513–9.

41. Béliveau C, Potvin C, Trudel J, Asselin A, Bellemare G. Cloning, sequencing, and expression in Escherichia coli of a Streptococcus faecalis autolysin. J Bacteriol. 1991;173(18):5619–23.

42. Abranches J, Tijerina P, Avilés-Reyes A, Gaca AO, Kajfasz JK, Lemos JA. The cell wall-targeting antibiotic stimulon of Enterococcus faecalis. PLoS One. 2014;8(6):e64875.

43. Darnell RL, Knottenbelt MK, Todd Rose FO, Monk IR, Stinear TP, Cook GM. Genomewide Profiling of the Enterococcus faecalis Transcriptional Response to Teixobactin Reveals CroRS as an Essential Regulator of Antimicrobial Tolerance. mSphere. 2019;4(3).

44. Kellogg SL, Kristich CJ. Functional Dissection of the CroRS Two-Component System Required for Resistance to Cell Wall Stressors in Enterococcus faecalis. J Bacteriol. 2016;198(8):1326–36.

45. Muller C, Massier S, Le Breton Y, Rincé A. The role of the CroR response regulator in resistance of Enterococcus faecalis to D-cycloserine is defined using an inducible receiver domain. Mol Microbiol. 2018;107(3):416–27.

46. Opsata M, Nes IF, Holo H. Class IIa bacteriocin resistance in Enterococcus faecalis V583: the mannose PTS operon mediates global transcriptional responses. BMC Microbiol. 2010;10:224.

47. Ran S, Liu B, Jiang W, Sun Z, Liang J. Transcriptome analysis of Enterococcus faecalis in response to alkaline stress. Front Microbiol. 2015;6:795.

48. Gaca AO, Lemos JA. Adaptation to Adversity: the Intermingling of Stress Tolerance and Pathogenesis in Enterococci. Microbiol Mol Biol Rev. 2019;83(3).

49. Flahaut S, Hartke A, Giard JC, Auffray Y. Alkaline stress response in Enterococcus faecalis: adaptation, cross-protection, and changes in protein synthesis. Appl Environ Microbiol. 1997;63(2):812–4.

50. Ran S, He Z, Liang J. Survival of Enterococcus faecalis during alkaline stress: changes in morphology, ultrastructure, physiochemical properties of the cell wall and specific gene transcripts. Arch Oral Biol. 2013;58(11):1667–76.

51. Fitzgerald BA, Wadud A, Slimak Z, Slonczewski JL. Enterococcus faecalis OG1RF Evolution at Low pH Selects Fusidate-Sensitive Mutants in Elongation Factor G and at High pH Selects Defects in Phosphate Transport. Appl Environ Microbiol. 2023;89(6):e0046623.

52. Arias CA, Panesso D, McGrath DM, Qin X, Mojica MF, Miller C, et al. Genetic basis for in vivo daptomycin resistance in enterococci. N Engl J Med. 2011;365(10):892–900.

53. Reyes J, Panesso D, Tran TT, Mishra NN, Cruz MR, Munita JM, et al. A liaR deletion restores susceptibility to daptomycin and antimicrobial peptides in multidrug-resistant Enterococcus faecalis. J Infect Dis. 2015;211(8):1317–25.

54. Comenge Y, Quintiliani R, Li L, Dubost L, Brouard JP, Hugonnet JE, et al. The CroRS two-component regulatory system is required for intrinsic beta-lactam resistance in Enterococcus faecalis. J Bacteriol. 2003;185(24):7184–92.

55. Paradis-Bleau C, Kritikos G, Orlova K, Typas A, Bernhardt TG. A genome-wide screen for bacterial envelope biogenesis mutants identifies a novel factor involved in cell wall precursor metabolism. PLoS Genet. 2014;10(1):e1004056.

56. Djorić D, Kristich CJ. Oxidative stress enhances cephalosporin resistance of Enterococcus faecalis through activation of a two-component signaling system. Antimicrob Agents Chemother. 2015;59(1):159–69.

57. Wang QQ, Zhang CF, Chu CH, Zhu XF. Prevalence of Enterococcus faecalis in saliva and filled root canals of teeth associated with apical periodontitis. Int J Oral Sci. 2012;4(1):19–23.

58. Pinto KP, Barbosa AFA, Silva EJNL, Santos APP, Sassone LM. What Is the Microbial Profile in Persistent Endodontic Infections? A Scoping Review. J Endod. 2023;49(7):786–98.e7.

59. Lindgren JK, Thomas VC, Olson ME, Chaudhari SS, Nuxoll AS, Schaeffer CR, et al. Arginine deiminase in Staphylococcus epidermidis functions to augment biofilm maturation through pH homeostasis. J Bacteriol. 2014;196(12):2277–89.

60. Thurlow LR, Joshi GS, Clark JR, Spontak JS, Neely CJ, Maile R, et al. Functional modularity of the arginine catabolic mobile element contributes to the success of USA300 methicillin-resistant Staphylococcus aureus. Cell Host Microbe. 2013;13(1):100–7.

61. Tian J, Utter DR, Cen L, Dong PT, Shi W, Bor B, et al. Acquisition of the arginine deiminase system benefits epiparasitic Saccharibacteria and their host bacteria in a mammalian niche environment. Proc Natl Acad Sci U S A. 2022;119(2).

62. Ladjouzi R, Bizzini A, van Schaik W, Zhang X, Rincé A, Benachour A, et al. Loss of Antibiotic Tolerance in Sod-Deficient Mutants Is Dependent on the Energy Source and Arginine Catabolism in Enterococci. J Bacteriol. 2015;197(20):3283–93.

63. Teng F, Singh KV, Bourgogne A, Zeng J, Murray BE. Further characterization of the epa gene cluster and Epa polysaccharides of Enterococcus faecalis. Infect Immun. 2009;77(9):3759–67.

64. Guerardel Y, Sadovskaya I, Maes E, Furlan S, Chapot-Chartier MP, Mesnage S, et al. Complete Structure of the Enterococcal Polysaccharide Antigen (EPA) of Vancomycin-Resistant Enterococcus faecalis V583 Reveals that EPA Decorations Are Teichoic Acids Covalently Linked to a Rhamnopolysaccharide Backbone. mBio. 2020;11(2).

65. Andrews S. FastQC: A quality control tool for high throughput sequence data: Babraham Bioinformatics; 2010 [Available from: https://www.bioinformatics.babraham.ac.uk/projects/fastqc/.

66. Bolger AM, Lohse M, Usadel B. Trimmomatic: a flexible trimmer for Illumina sequence data. Bioinformatics. 2014;30(15):2114–20.

67. McClure R, Balasubramanian D, Sun Y, Bobrovskyy M, Sumby P, Genco CA, et al. Computational analysis of bacterial RNA-Seq data. Nucleic Acids Res. 2013;41(14):e140.

68. Tjaden B. De novo assembly of bacterial transcriptomes from RNA-seq data. Genome Biol. 2015;16:1.

69. Schindelin J, Arganda-Carreras I, Frise E, Kaynig V, Longair M, Pietzsch T, et al. Fiji: an open-source platform for biological-image analysis. Nat Methods. 2012; 9(7):676–82.

